# Industrial chicory genome gives insights into the molecular timetable of anther development and male sterility

**DOI:** 10.1101/2023.02.08.527727

**Authors:** Evelien Waegneer, Stephane Rombauts, Joost Baert, Nicolas Dauchot, Annick De Keyser, Tom Eeckhaut, Annelies Haegeman, Chang Liu, Olivier Maudoux, Christine Notté, Ariane Staelens, Jeroen Van der Veken, Katrijn Van Laere, Tom Ruttink

**Affiliations:** ILVO, Flanders Research Institute for Agriculture, Fisheries and Food, Plant Sciences Unit, Melle, Belgium; Laboratory for Plant Genetics and Crop Improvement, Division of Crop Biotechnics, Department of Biosystems, Katholieke Universiteit Leuven, Leuven, Belgium; Department of Plant Biotechnology and Bioinformatics, Ghent University, 9052 Ghent, Belgium; Center for Plant Systems Biology, VIB, 9052 Ghent, Belgium; Unit of Cellular and Molecular Plant Biology, UNamur, 5000 Namur, Belgium; Department of Epigenetics, Institute of Biology, University of Hohenheim, 70599 Stuttgart, Germany; Chicoline, a division of Cosucra Groupe Warcoing S.A., 7740 Warcoing, Belgium

## Abstract

Industrial chicory (*Cichorium intybus* var. *sativum*) is a biannual crop mostly cultivated for extraction of inulin, a fructose polymer used as a dietary fiber. F_1_ hybrid breeding is a promising breeding strategy in chicory but crucially relies on stable self-incompatibility. Here, we report the assembly and annotation of a new industrial chicory reference genome. Additionally, we performed RNA-Seq on subsequent stages of flower bud development of a fertile line and two cytoplasmic male sterile (CMS) clones. Comparison of fertile and CMS flower bud transcriptomes combined with morphological microscopic analysis of anthers, provided a molecular understanding of anther development and identified key genes in a range of underlying processes, including tapetum development, sink establishment, pollen wall development and anther dehiscence. We also described the role of phytohormones in the regulation of these processes under normal fertile flower bud development. In parallel, we evaluated which processes are disturbed in CMS clones and could contribute to the male sterile phenotype. Taken together, this study provides a state-of-the-art industrial chicory reference genome, an annotated and curated candidate gene set related to anther development and male sterility as well as a detailed molecular timetable of flower bud development in fertile and CMS lines.

## INTRODUCTION

Chicory (*Cichorium intybus L.*) is a biannual crop belonging to the *Asteraceae* family. The species is cultivated as a leafy vegetable (e.g., witloof) or as a root crop (e.g., industrial chicory) because of its high content of inulin, a fructose polymer that is used as a prebiotic soluble dietary fiber (Shoaib, Shehzad et al. 2016, Aldahak, Salem et al. 2021). One of the breeding strategies used in chicory is hybrid breeding, where two compatible homozygous lines are crossed to obtain high yielding heterotic F_1_ offspring. Mass production of hybrid seed requires male sterile lines to prevent self-pollination, as male sterility inherently eliminates the need for manual emasculation or chemical treatment to inhibit pollen growth. Cytoplasmic male sterility (CMS) has been described in many crops (reviewed by Bohra, Jha et al. (2016)). The male sterility phenotype in CMS is maternally inheritable and often caused by altered mitochondrial genes. The CMS phenotype can be suppressed by nuclear *Restorer-of-fertility* genes, indicating that defective communication between mitochondria and nucleus might be at the basis. CMS can occur either spontaneously, or can be induced via intra- or interspecific protoplast fusion or mutagenesis. Elucidation of the mechanisms causing CMS requires knowledge on two aspects. First, a comprehensive understanding of normal flower bud development leading to fertile and mature pollen, including a detailed description of the underlying morphological, physiological, and molecular processes; and second, an inventory of the processes that are affected in CMS lines to delineate the possible causes and downstream effects that prevent pollen development and maturation.

Driven by the importance of CMS for hybrid breeding in agricultural crops, considerable efforts to elucidate the underlying mechanisms have unveiled an array of causative genes and morphological symptoms across various species (reviewed by Hanson and Bentolila (2004), Chase (2007), Chen and Liu (2014) and Kim and Zhang (2018). Four models for possible CMS mechanisms are supported by currently available evidence; the cytotoxicity model, the energy deficiency model, the aberrant programmed cell death (PCD) model and the retrograde regulation model (Chen and Liu 2014). The cytotoxicity model assumes that a CMS-causing protein is cytotoxic for the cell, thereby killing sporophytic and gametophytic tissues in the anther. The energy deficiency model proposes that male gametophytic development requires a high amount of energy and CMS mitochondria fail to meet these energy requirements in stamen, resulting in abortion of pollen. The aberrant PCD model predicts that male sterility is the result of disturbed timing of PCD of the tapetum, a tissue in the anther wall with an important role in feeding developing pollen. Lastly, the retrograde regulation model suggests that mitochondrial proteins alter nuclear gene expression via retrograde signals, which in turn cause the defects that result in male sterility. Previous research has shown that CMS involves complex processes, and therefore it is likely that most CMS phenotypes can be caused by combinations of these four models. In chicory, CMS studies have been limited to a morphological study of line CMS524 (Habarugira, Hendriks et al. 2015) and identification of the causative mitochondrial defect in a CMS cybrid (Rambaud, Bellamy et al. 1997, Varotto, Nenz et al. 2001).

While most studies on chicory fertile flower bud development have focused on morphology, molecular studies are limited to flower initiation (Perilleux, Pieltain et al. 2013, Mathieu, Perilleux et al. 2020), and a detailed temporal description of molecular events during flower bud development is lacking. Starting from flower initiation, flower bud development to anthesis takes approximately 15 days in chicory, and consists of a differentiation phase and a pollen maturation phase (Habarugira, Hendriks et al. 2015). The differentiation phase consists of the formation of floral organs and differentiation of the anther walls into epidermis, endothecium, middle layer, and tapetum cells, surrounding the developing pollen (Habarugira, Hendriks et al. 2015). The subsequent pollen maturation phase is marked by degeneration of the middle layer, tapetum, septum, and stomium to facilitate pollen release.

Currently, a detailed description of the molecular events driving flower bud development in chicory is lacking, yet is critical to understand the molecular mechanisms leading to CMS. Here, we studied two chicory CMS clones; CMS36 and CMS30. These CMS clones were developed by crossing a wild chicory genotype with an inbred line of industrial chicory (Van der Veken, Vandermoere et al. 2018). In subsequent backcrosses with industrial chicory progressively fewer pollen was produced until no pollen was present in the third and fourth backcross generation. While the cause and mechanism of male sterility is unknown in these clones, CMS may result from the incompatibility between wild chicory cytoplasm and industrial chicory nuclei. However, in both CMS clones fertility could be restored at elevated temperature (Van der Veken, Vandermoere et al. 2018). The response time between heat shock and fertility restoration was three weeks, suggesting that early stage flower buds (size between 2 mm and 4 mm) are most responsive to fertility restoration. CMS30 male sterility was more robust as only few pollen were produced after a heat shock, whereas in CMS36 full fertility restoration was observed (Van der Veken, Vandermoere et al. 2018).

With the aim to gain insight in the molecular regulation of flower bud development in fertile chicory and to uncover transcriptome changes associated with CMS, we first created a chromosome-scale reference genome sequence for *C. intybus* var. *sativum* (industrial chicory) and identified a functionally annotated gene set. Using that gene set as reference, we performed RNA-Seq transcriptome sequencing at eight subsequent stages of flower bud development in a fertile line and two CMS clones, CMS36 and CMS30. The temporal expression profiles of thousands of differentially expressed genes were then clustered to delineate co-expressed gene sets that, in turn, were used to dissect the temporal sequence of events during flower bud development using the gene functional annotations per co-expression cluster. Transient activation of regulatory networks, hormonal signaling pathways, and biochemical or metabolic pathways captured by the temporal expression clusters were anchored to cellular events by morphological analysis via light microscopy of developing flower buds. Thus, we give a comprehensive overview of the molecular processes during anther development, including ethylene, jasmonate, auxin, brassinosteroid and gibberellin signaling and response, regulation of sporopollenin biosynthesis, energy, phenylpropanoid and isoprenoid metabolism, tapetum development and degradation, pollen development and maturation, and anther dehiscence. Additionally, we analyzed which of these developmental, biochemical, metabolic, and hormonal regulatory pathways were disturbed in clones CMS36 and CMS30, and evaluate those molecular signatures in the context of the four CMS models known from other species. Taken together, these studies provide a reference genome sequence with functionally annotated gene sets as state-of-the-art genomics resource for chicory, validated by manual curation of hundreds of candidate genes assigned to a broad range of genetic networks, thus paving the way for future structural genome biology and comparative genomics studies. They further provide a detailed timetable of molecular events that can be used as reference framework to profile the molecular phenotypes of a range of additional CMS clones to investigate which other modes of action may cause CMS in chicory, and identified key regulatory components that may be targeted to modulate CMS phenotypes.

## RESULTS

### Cichorium intybus genome assembly and gene annotation

An eighth generation inbred line (L8001) of *C. intybus* var. *sativum* (industrial chicory) was used for Whole-Genome Sequencing (WGS) and assembly. Genomic read data were created with Illumina HiSeq2500 (WGS, PE-100, 488M read-pairs, coverage 40x) NextSeq (WGS, PE-100, 394M read-pairs, 33x), MiSeq (WGS, PE-295, 24M read-pairs, 6x), PacBio RS II (1.38 M reads; range 1001 bp to 38,4 kb; 7.8 Gb; read-N50 8.3 kb; genome coverage ∼7x) and Oxford Nanopore Technologies (ONT) MinION (2.99 M reads; range 100 bp to 694 kb; 16.8 Gb; read-N50 10.7 kb; genome coverage ∼15x). In short, the primary assembly was generated with Flye (Kolmogorov, Yuan et al. 2019), using the ONT reads. Long read polishing was performed with Racon using the ONT and PacBio reads combined, and further polishing was performed with Pilon using Illumina WGS short reads. Hi-C proximity ligation libraries (PE-120, 46M reads) were used to construct a contact probability map, and scaffolds were ordered and oriented with 3D-DNA/Juicer. The Hi-C data allowed for the scaffolding into chromosome arms, but a linkage map was needed to confirm, join, and order these into the final 9 chromosomes with a total chromosome-anchored length of 877 Mb (leaving about 28.6 Mb unanchored) (Supplemental Table 1). Similarly to previous publications in lettuce and chicory, we observe triplication patterns across the genome (Figure 1) (Reyes-Chin-Wo, Wang et al. 2017, Fan, Wang et al. 2022).

**Figure 1.**
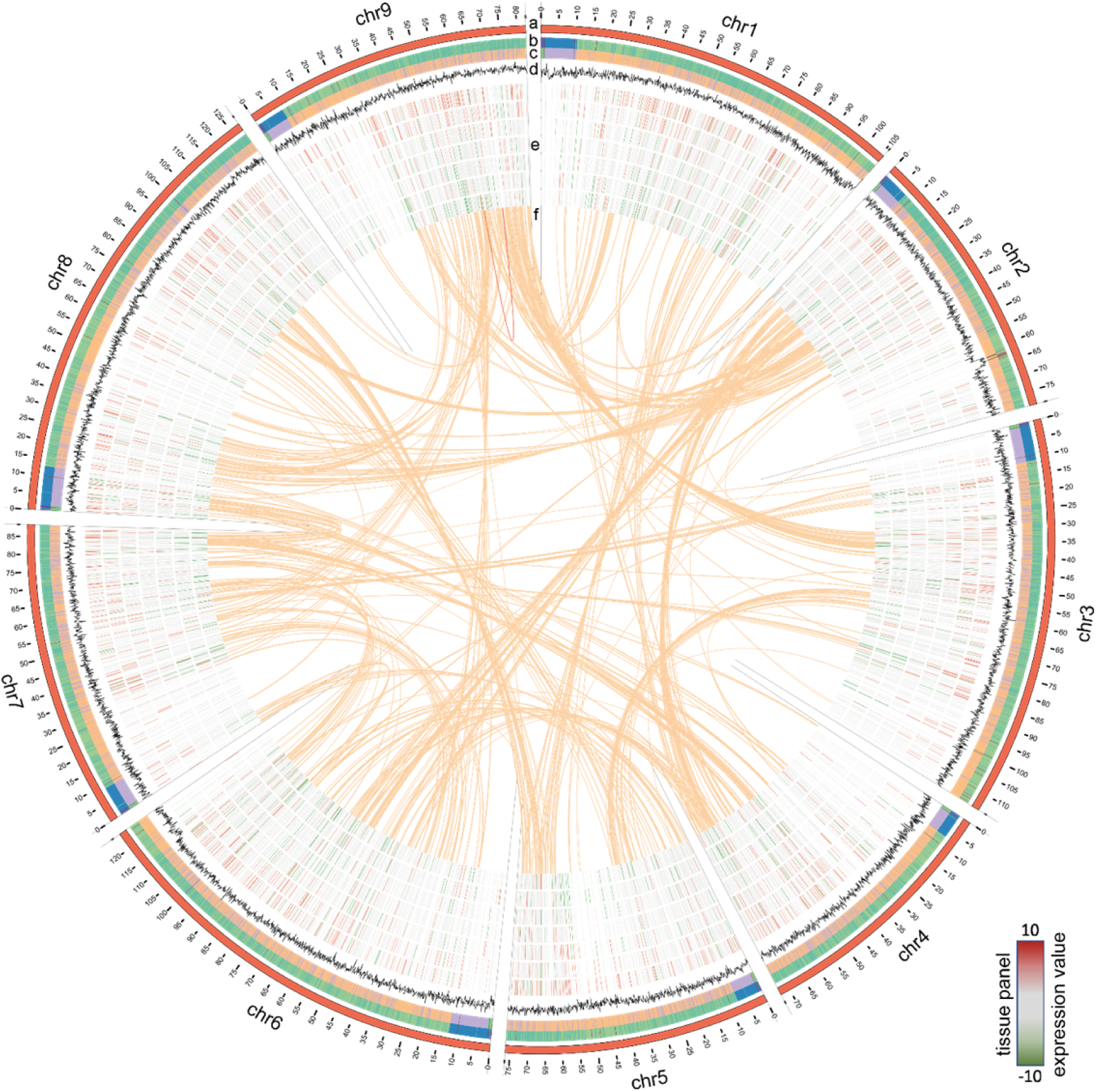
Circos plot representing the *C. intybus* L8001 genome. a) chromosome-scale assembly (in Mbp), b) heatmap for gene density, c) heatmap for repeat density, d) GC%, e) expression profile of selected genes (outer-inner: Anther, Ligule, Ovule, Petal, Leaf, Seedling, Root, RPKM values ranging from [-10..10] with low expression indicated in green and high expression in red), f) synteny blocks based on genome duplications as reported by i-ADHoRe.

Augustus was trained prior to gene prediction using the eudicots gene set from BUSCO. *De novo* repeat detection was performed on the final assembly and the genome was repeat-masked prior to gene prediction (Figure 1). Augustus predicted 53386 genes (Figure 1) using hints from RNA-Seq of the flower bud developmental series and a small tissue panel (Figure 1) and proteins from reference proteomes (*Arabidopsis,* tomato, lettuce) as biological evidence (Supplemental Table 1). BUSCO genome completeness analysis using the eudicots reference set (n=2326 genes) resulted in C:92.7% [S:88.9%, D:3.8%], F:1.2%, M:6.1%, thus only missing 140 genes from the whole BUSCO reference set. Manual curation of predicted genes putatively involved in hormonal, metabolic, and regulatory pathways was done via ORCAE, combined with orthology identification via detailed gene family analyses, protein alignment, phylogenetic analyses, and clade classification (Supplemental Data Set 2 lists all genes, functional description, assignment per pathway as described in Figures 2, 4 and 5, and normalized expression values during flower bud development). The reference genome sequence, all annotated gene models, functional description, and expression data in a small tissue panel is publicly available via ORCAE (https://bioinformatics.psb.ugent.be/orcae/overview/Cicin).

**Figure 2.**
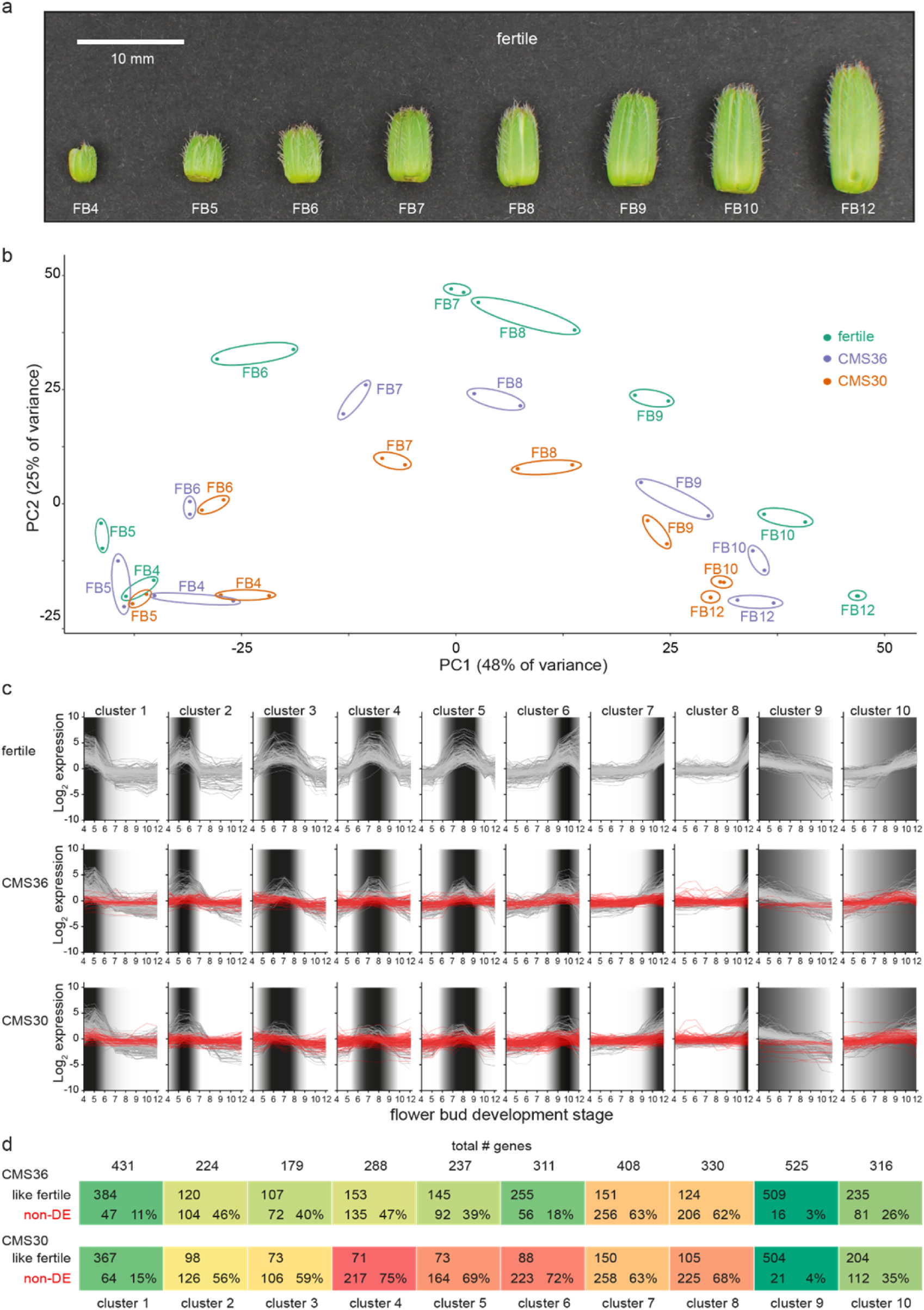
Global transcriptome analysis of flower bud development in fertile line K1337 and male sterile CMS clones CMS36 and CMS30 of industrial chicory (*Cichorium intybus* var. *sativum*). a) Developmental series of flower buds (FB) at increasing size (4-12mm) sampled for microscopy and RNA-Seq. b) Principal component analysis on 30971 transcript expression profiles. c) Ten clusters of differential expression (DE) profiles identified within the> 4-fold-change gene set of the fertile line. Dark shaded areas indicate the periods of high relative expression level. d) Comparison of expression profiles between fertile line and CMS lines, either having a similar expression profile to the fertile line (like fertile), or DE in fertile line but not DE across the developmental series in the corresponding CMS line (non-DE).

### Temporal transcriptome profiling during flower bud development

The chicory flower bud developmental process was reconstructed by sampling a series of flower buds of increasing sizes (4-12 mm, see Materials and Methods), reflecting eight progressive stages of flower bud development (referred to as FB4 – FB12), in a fertile line and the two CMS clones, CMS30 and CMS36 (Figure 2a). These developmental sample series were subjected to light microscopy to map morphological development, and to transcriptome profiling to yield temporal expression profiles for 30971 out of 53386 predicted genes (58%), with detectable expression levels across all samples (see Materials and Methods). Principal component analysis (PCA) on the 30971 transcript expression profiles (Figure 2b) revealed that subsequent stages of flower bud development (FB4-12) of all three lines are resolved along the first principal component axis, suggesting that the majority of the expression variance in the dataset (PC1, 48%) reflects temporal changes in the transcriptome throughout flower bud development. Differences between transcriptomes of fertile and CMS clones are resolved on the second principal component axis (PC2, 25%) (Figure 2b). The PCA showed that in early stages (FB4-5) samples from fertile buds cluster closely together with samples from CMS buds, suggesting little initial transcriptional differences between fertile and CMS flower buds. From FB6 onwards, PCA analysis showed that the transcriptional difference between fertile and CMS buds progressively increased (Figure 2b). Accordingly, microscopic analysis of morphological development revealed that CMS anther morphology initially does not markedly differ from fertile anther morphology, except for slight differences in tapetum degradation and pollen morphology in FB4-7 (CMS36) and FB4-6 (CMS30) (Figure 3a-c,i-k,q-s). In later stages (FB8/FB9), CMS clones are characterized by early endothecium lignification (FB6) and anther dehiscence in FB8/FB9 (CMS36/CMS30), while CMS30 additionally displays inwards collapse of the anther and early pollen degradation (Figure 3f,h,k,m,s,v).

**Figure 3.**
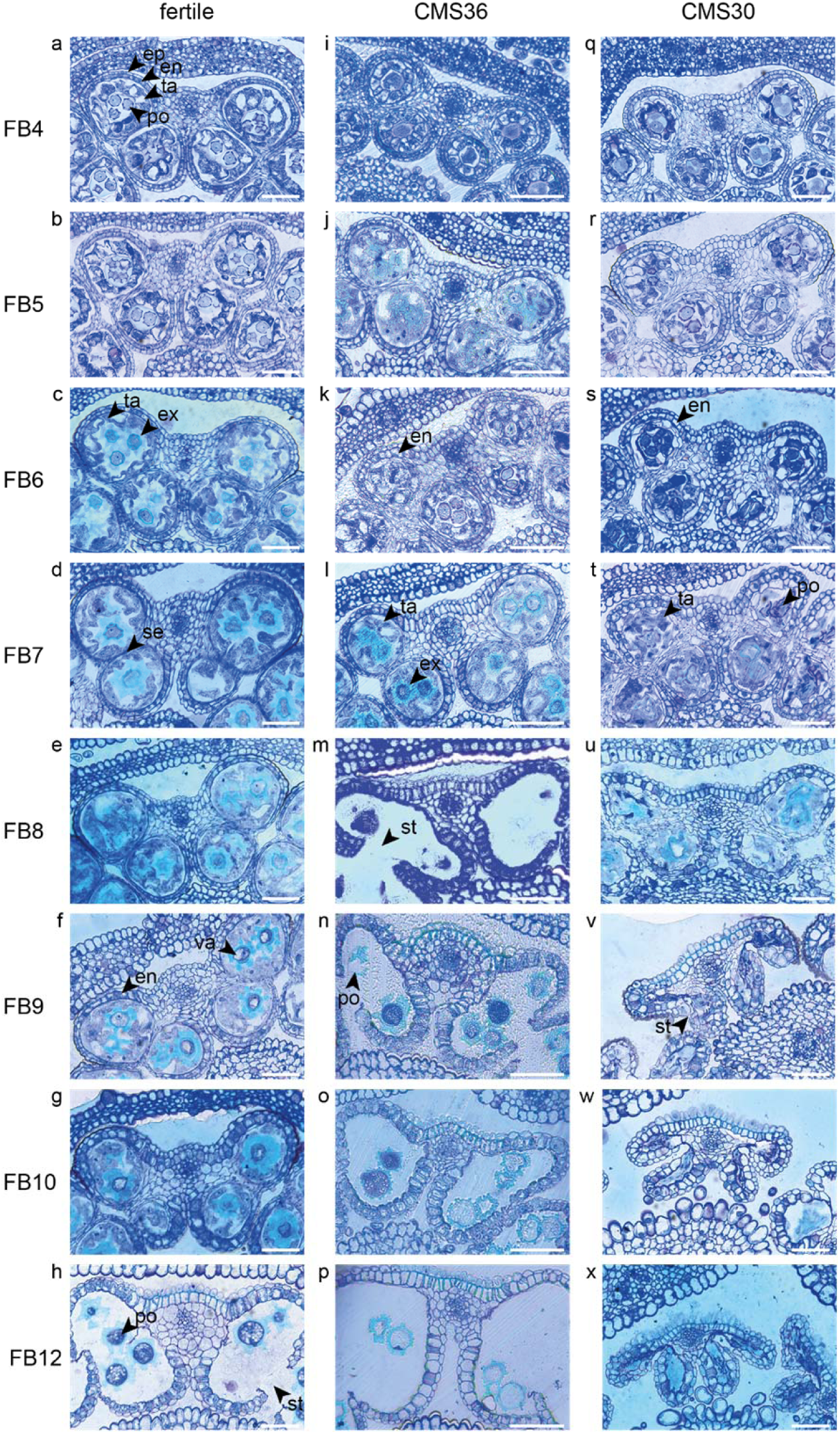
Morphological analysis and comparison of anthers of fertile line L8018 and male sterile CMS clones CMS36 and CMS30 of industrial chicory (*Cichorium intybus* var. *sativum*). a) fertile 4mm flower bud (FB) stage with epidermis (ep), endothecium (en), tapetum (ta) and pollen (po). b) fertile 5mm FB stage. c) fertile 6mm FB stage with degraded tapetum cell walls (ta) and developing exine walls of pollen (ex). d) fertile 7mm FB stage with the start of septum degradation (se). e) fertile 8mm FB stage. f) fertile 9mm FB stage with the start of endothecium expansion and lignification (en) and pollen vacuolization (va). g) fertile 10mm FB stage. h) fertile 12mm FB stage with stomium opening (st) and pollen starch bodies (po). i) CMS36 4mm FB stage. j) CMS36 5mm FB stage. k) CMS36 6mm FB stage with the start of endothecium expansion and lignification (en). l) CMS36 7mm FB stage with degraded tapetum cell walls (ta) and start of pollen exine wall formation (po). m) CMS36 8mm FB stage with stomium opening (st). n) CMS36 9mm FB stage with start of pollen degradation (po). o) CMS36 10mm FB stage. p) CMS36 12mm FB stage. q) CMS30 4mm FB stage. r) CMS30 5mm FB stage. s) CMS30 6mm FB stage with start of endothecium expansion and lignification (en). t) CMS30 7mm FB stage with degraded tapetum cell walls (ta) and start of pollen degradation (po). u) CMS30 8mm FB stage. v) CMS30 9mm FB stage with stomium opening (st) and inwards collapse of the anther. w) CMS30 10mm FB stage. x) CMS30 12mm FB stage.

Next, we performed likelihood ratio tests (LRT) on the fertile line gene expression profiles, and identified 21247 statistically significant (p<0.05) differentially expressed genes (DEG) within the fertile flower bud developmental series. This indicates that the vast majority of genes expressed in flower buds (69% of 30971 expressed genes) showed dynamic expression during flower bud development. Of these, 8039 genes (25%) displayed at least a 2-fold-change in expression, while 4067 genes displayed a >4-fold-change in expression (13%, from hereon called the FC4 gene set). Next, we looked at expression levels of FC4 gene set in the tissue panel (anther, ligule, ovule, petal, leaf, root and seedling) to assess flower bud development specific expression (Figure 1, Supplemental Data Set 1). As expected, higher expression of these genes was observed in flower tissues (anther, ligule, ovule and petal), whereas in leaf, seedling and root tissues these genes were less expressed (Figure 1e).

The expression profiles of the FC4 gene set within the fertile time series were then clustered to investigate the temporal sequence of events, and to delineate subsequent biological processes during flower bud development. Of the 4067 FC4 genes, 3279 genes (80%) were divided into 10 clusters; one cluster with 316 gradually up-regulated genes and one cluster with 525 gradually downregulated genes, and eight clusters containing genes transiently expressed in specific FB stages, thus revealing overall very high levels of co-expression (Figure 2c, Supplemental Data Set 1). Per cluster, genes with similar transcriptional upregulation in the fertile line and both CMS clones were separated from genes with little or no transcriptional upregulation in CMS36 and/or CMS30 (Figure 2c-d). Consistent with the global PCA analysis, a relatively small fraction of genes in clusters transiently expressed during early stages was not upregulated in CMS clones. The fraction of genes without transcriptional upregulation in CMS clones became more pronounced in clusters expressed during later stages of flower bud development. The fraction of genes without transcriptional upregulation in CMS30 was also consistently higher than that fraction of genes in CMS36 (Figure 2d), which could correspond to the severity of morphological differences observed by microscopy (Figure 3), or the different stability of sterility between the two CMS clones. This means that in general, processes that normally occur in fertile flowers are suppressed or absent in the CMS clones, and the associated gene expression in those cell types or organs is also not activated normally at the appropriate time. Thus, the comparative analysis of fertile versus CMS clones allows to uncouple the developmental processes that occur simultaneously in whole flower buds and this, in turn, adds resolution to delineate co-expressed genes and disentangle molecular processes.

### Dissecting molecular processes during flower bud development

In the following, we performed a targeted analysis based on the functional description of genes, assigned genes to known regulatory pathways or metabolic processes based on their functional annotation, combined with their temporal expression dynamics, to describe the activity of a range of biological processes during subsequent stages of chicory flower bud development. To further strengthen the high quality of the reference genome sequence and gene annotation, we manually curated the gene model structure of all candidate genes described in detail below (for further details, see Supplemental Data Set 2) and eliminated pseudogenes to ensure accurate expression profiling and functional annotation, and pave the way for future functional studies on individual genes.

### Early tapetum development

The tapetum is a cell layer in the anther that mainly functions as feeding tissue for developing pollen. An important function of the tapetum layer is synthesis of sporopollenin, a pollen wall precursor (Wang, Guo et al. 2018). The developmental timing of tapetum development and degradation is important to generate viable pollen and tapetum defects lead to male sterility. To investigate whether male sterility in the clones CMS36 and CMS30 involves defects in tapetum development, functioning, or degradation, we analyzed expression profiles of important regulators of tapetum development, as well as genes involved in tapetum function. For instance, in a genetic pathway proposed by Li, Xue et al. (2017), *DYT1* and *TDF1* are transcription factors regulating tapetum development, while *AMS* and *MS188* regulate sporopollenin synthesis, tapetum degradation and pollen wall development. In *Arabidopsis, AMS* regulates *TEK* expression, important for sexine and nexine formation in the two layers of the outer pollen wall (exine) (Li, Xue et al. 2017). *MS1* is a transcription factor involved in late tapetum development and pollen wall formation in *Arabidopsis* (Li, Xue et al. 2017, Wang, Guo et al. 2018). Here, light microscopic analysis revealed that degradation of tapetum cell walls, an early hallmark of tapetum degradation, takes place around FB6 in the fertile line (Figure 3c). Accordingly, microscopic analysis revealed that CMS36 and CMS30 display a global pattern of tapetum degradation that is similar to the fertile line, except that the cell walls are still intact at FB6 in CMS36 and CMS30 and degradation only starts at FB7 (Figure 3k-l,s-t), suggesting a slight delay of tapetum developmental timing. As expected, key transcription factors regulating tapetum development (*CiTDF1*, *CiMS188*, and *CiTEK*) were highly expressed just prior to tapetum degradation (FB4-FB5) in the fertile line (Figure 4a). Two putative *CiMS1* genes were highly expressed during early stages (FB4, FB5 and FB6) (Figure 4a). Furthermore, another eight genes involved in sporopollenin biosynthesis were all highly expressed at stages FB4 and FB5, and were strongly down-regulated at FB7, consistent with degradation of the tapetum layer (Figure 4a). In both CMS clones, *CiTDF1*, *CiMS188*, *CiTEK*, *CiMS1*, and sporopollenin biosynthesis genes showed expression profiles similar to the fertile line. However, in both CMS clones, high expression of these genes was generally maintained until FB7 instead of FB6 (Figure 4a), consistent with a slight delay of tapetum developmental timing (Figure 3k-l,s-t). Additionally, we searched for core marker genes for developmental programmed cell death (dPCD) and identified five genes encoding aspartic protease (Olvera-Carrillo, Van Bel et al. 2015). These genes were highly upregulated from FB6 until FB9 in fertile lines, coinciding with tapetum cell wall degradation (Figure 4a). In CMS36, upregulation of four out of five genes was delayed until FB7 and maintained until FB8, while in CMS30, upregulation of these four was similar to the fertile line but less strong (Figure 4a). The fifth aspartic protease did not show any upregulation in CMS36 or CMS30 (Figure 4a). Taken together, this confirms that tapetum development and degradation is delayed in CMS clones, but not severely disturbed.

**Figure 4.**
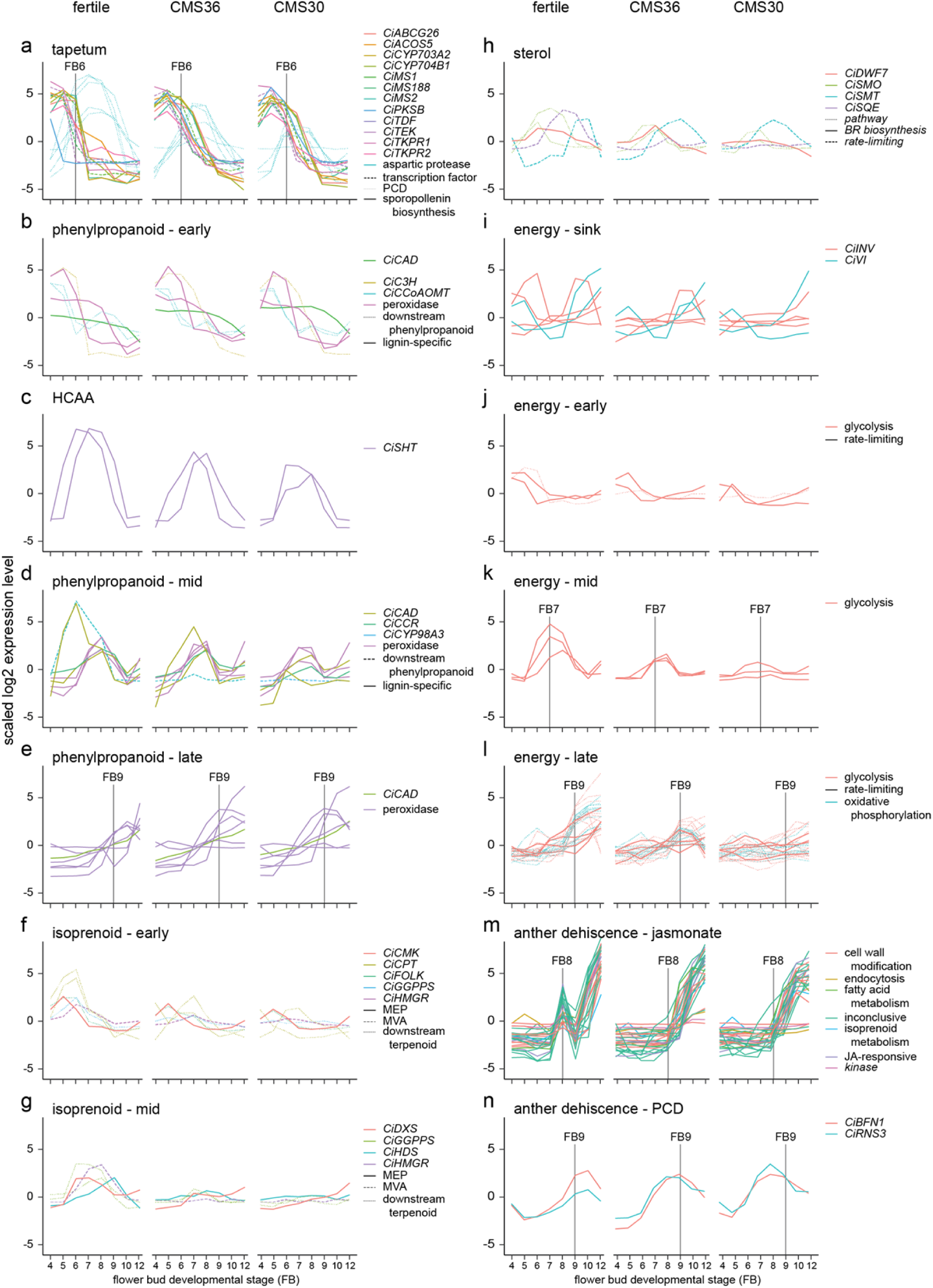
Expression profiles of genes involved in specific pathways important for anther processes of fertile line K1337 and male sterile CMS clones CMS36 and CMS30 of industrial chicory (*Cichorium intybus* var. *sativum*). x-axis shows flower bud developmental stages, ranging from 4mm (FB4) to 12mm (FB12). y-axis shows scaled log2 Reads Per Kilobase transcript per Million reads mapped (RPKM) values. Individual genes are shown with different colors, while pathway grouping is shown by different line types. Gene identifiers per subgroup are listed in Supplemental Data Set 2. For information on ortholog assignment, see Materials and Methods.

### Phenylpropanoid metabolism

The phenylpropanoid pathway is part of secondary metabolism and produces a variety of phenolic compounds in plants including precursors for lignin, flavonoids and suberin, a component of the pollen wall (Ma and Constabel 2019). These compounds have many functions during flower bud development, like providing building blocks for the pollen wall and several lignification processes (Battat, Eitan et al. 2019). Lignin biosynthesis and breakdown plays a key role during anther development, most notably during endothecium lignification and anther dehiscence. In the fertile line, genes involved in phenylpropanoid metabolism and monolignol biosynthesis show three different expression profiles during flower bud development (Figure 4b-e). For instance, during FB4-6, the upregulated genes encode enzymes such as *CINNAMYL ALCOHOL DEHYDROGENASE* (*CiCAD)* and *CAFFEOYL-COA O-METHYLTRANSFERASE (CiCCoAOMT)* acting in the phenylpropanoid pathway that produces monolignols (Figure 4b). In FB7-9, the majority of upregulated genes encode monolignol biosynthesis enzymes (*CiCAD* and *CINNAMOYL-COA REDUCTASE (CiCCR))* or extracellular peroxidases (Figure 4d). In FB9-12, the vast majority of upregulated genes encode extracellular peroxidases (Figure 4e), putatively involved in monolignol polymerization at the cell wall (Xie, Zhang et al. 2018). This suggests that biosynthesis of precursors for flavonols and monolignols takes place until FB9, whereas during stages FB9-12 lignin polymerization becomes important. Accordingly, microscopic analysis revealed that endothecium lignification started at FB9 in the fertile line. In CMS clones, genes upregulated during FB4-5 and FB6-9 show expression dynamics similar to the fertile line (Figure 4b-d). In late stages (FB9-12), one peroxidase is upregulated earlier in CMS clones (FB7) compared to the fertile line (FB9), which is consistent with earlier endothecium lignification in CMS clones. Two other peroxidases are not upregulated in CMS clones in FB12, while the fertile line does show upregulation in FB12 (Figure 4e). Taken together, the changed expression patterns of mostly peroxidases could reflect the earlier endothecium lignification, and earlier anther dehiscence in CMS clones.

The phenylpropanoid pathway also results in biosynthesis of other phenolic compounds such as hydroxycinnamic acid amides (HCAAs) (Figure 4c). Specific conjugates from HCAAs, i.e., hydroxycinnamoyl spermidines, are produced in the tapetum and accumulate in the *Arabidopsis* pollen coat (Fellenberg, Bottcher et al. 2009), and play a key role in pollen development. In sunflower, two *SPERMIDINE HYDROXYCINNAMOYL TRANSFERASE (SHT)* genes are hub genes regulating the flux from the phenylpropanoid pathway into the HCAA pathway during pollen development (Li, Yaermaimaiti et al. 2021). The chicory orthologs of these genes (*CiSHT*) show peak expression at FB6-8 in fertile flower buds (Figure 4c). In CMS36 and CMS30, *CiSHT* is reduced, with stronger reduction in CMS30 compared to CMS36 (Figure 4c). Together these results indicate that HCAA biosynthesis, important pollen wall components, take place during FB6-8, and is decreased in CMS clones, but not severely disturbed.

### Isoprenoid metabolism

Isoprenoid metabolism produces precursors for a variety of compounds with numerous functions, including plant hormones such as gibberellin, brassinosteroids, cytokinins and abscisic acid (Vranova, Coman et al. 2013). Additionally, isoprenoids are involved in photosynthetic processes, and many sterols in plants are produced via the isoprenoid metabolic pathway (Vranova, Coman et al. 2013). All isoprenoids are derived from the same five-carbon building block: isopentenyl diphosphate (IPP), which can be produced via two independent pathways in plants (Vranova, Coman et al. 2013, Tholl 2015). Some key differences between these two pathways distinguish their effect on male sterility. The mevalonate (MVA) pathway occurs in the cytosol and leads to the production of triterpenoids and sterols, whereas the 2-C-methyl-D-erythritol 4-phosphate (MEP) pathway takes place in plastids and is mostly important for production of carotenoids and chlorophyll side chains (Suzuki, Kamide et al. 2004, Vranova, Coman et al. 2013). Mutations in enzymes involved in the MVA pathway, but not the MEP pathway, have been linked to male sterility (Suzuki, Kamide et al. 2004, Okada, Kasahara et al. 2008). It is thought that male sterility may result from disturbed sterol composition in anthers, as an important sterol precursor, squalene, could rescue the male sterile phenotype in MVA pathway mutants (Suzuki, Kamide et al. 2004, Okada, Kasahara et al. 2008, Vranova, Coman et al. 2013).

We identified genes involved in both the MVA and MEP pathway and acting downstream in the terpenoid biosynthesis pathway. In the MVA pathway, we found two *HYDROXYMETHYLGLUTARYL-COA REDUCTASE* (*CiHMGR*) genes, encoding a rate-limiting enzyme in the biosynthesis pathway. One gene was upregulated in FB6, and the other in FB7-8 in the fertile line, while no upregulation was observed in CMS clones (Figure 4f,g). In the MEP pathway, we found *1-DEOXY-D-XYLULOSE 5-PHOSPHATE SYNTHASE* (*CiDXS*), a rate-liming enzyme*, 4-(CYTIDINE 5′-DIPHOSPHO)-2-C-METHYL-D-ERYTHRITOL KINASE* (*CiCMK*) and *4-HYDROXY-3-METHYLBUT-2-ENYL DIPHOSPHATE SYNTHASE* (*CiHDS*). *CiCMK* was upregulated at FB5 in the fertile line and both CMS clones (Figure 4f). *CiDXS* was upregulated at FB6-7 in the fertile line, while *CiHDS* was upregulated at FB9 (Figure 4g). In contrast, no upregulation of these genes was observed in the CMS clones (Figure 4f,g). Genes acting downstream in the terpenoid pathway included *CIS-PRENYLTRANSFERASE* (*CiCPT*), *FARNESOL KINASE* (*CiFOLK*) and *GERANYLGERANYL PYROPHOSPHATE SYNTHASE* (*CiGGPPS*). *CiCPT, CiFOLK*, and *CiGGPPS* were upregulated at FB5-6 in the fertile line, but in CMS clones, these genes were either not upregulated or upregulated at different stages (Figure 4f,g). Two other *CiGGPPS* genes were upregulated at FB6-8 in the fertile line, but no upregulation was observed in the CMS clones (Figure 4g). Taken together, these results suggest that both the MVA and MEP pathways, as well as the downstream terpenoid pathway are disrupted in CMS clones.

We identified four genes in the FC4 gene set involved in the sterol biosynthesis pathway: the rate-limiting enzyme SQUALENE EPOXIDASE (*CiSQE), C-24 STEROL METHYLTRANSFERASE (CiSMT)* and two *4 ALPHA-METHYL OXIDASEs* (*CiSMOs*). CMS clones displayed differences in gene expression of these genes compared to the fertile line, although these differences were less pronounced in CMS36 than in CMS30 (Figure 4h). Most noteworthy is the lack of upregulation in both CMS clones of the rate-limiting *CiSQE*, acting upstream in the sterol biosynthesis pathway (Shi, Gonzales et al. 1996, Carland, Fujioka et al. 2010, Doblas, Amorim-Silva et al. 2013), suggesting compromised sterol biosynthesis in both CMS clones (Figure 4h). In addition, we found a brassinosteroid biosynthesis gene (*CiDWF7)* acting downstream in the sterol biosynthesis pathway, which was slightly upregulated at FB6-8 in the fertile line, and had similar expression in CMS36, but not in CMS30 (Figure 4h). Disturbed sterol composition in anthers could also be a contributing factor to the male sterility phenotype, similar to previous observations in other plant species (Suzuki, Kamide et al. 2004, Okada, Kasahara et al. 2008, Vranova, Coman et al. 2013).

### Energy metabolism

Anther and pollen development are highly energy-demanding processes, and disturbances in carbohydrate allocation or energy metabolism can lead to male sterility (Chen and Liu 2014, Geng, Ye et al. 2018). An important factor controlling anther development is the increase of sink strength via cell wall invertase (INV) activity, which cleaves sucrose in glucose and fructose, compounds that can subsequently be transported into sink tissues (Zhang, Liang et al. 2010, Bihmidine, Hunter et al. 2013, Morey, Hirose et al. 2018, Chen, Qin et al. 2019, Yan, Wu et al. 2019). Therefore, we compared the expression profiles of genes involved in sink strength and glucose metabolism in the fertile line and CMS clones (Figure 4i-l).

In the fertile line, out of four *CiINV* genes, two genes were highly expressed in FB4-6, one gene in FB9-10, and one gene gradually increased expression during subsequent stages of flower bud development (Figure 4i), suggesting that the flower bud indeed acts as a strong sink during these stages. In addition to cell wall invertases, we also found two vacuolar invertases (*CiVI*) that were highly upregulated in both FB4-6 and FB9-12 (Figure 4i). Late stage upregulation of *CiVI* coincides with vacuolization in chicory pollen (Figure 3f-g, Figure 6). Such *CiVI* have been found to be highly upregulated during etiolated leaf growth in chicory and this is indicative of a growing sink tissue (Van den Ende, Michiels et al. 2002). In addition, as *VI* produces large amounts of fructose and glucose, an osmotic effect is generated that could be the driving force behind pollen vacuolization (Van den Ende, Michiels et al. 2002). Furthermore, genes involved in glucose metabolism (glycolysis and oxidative phosphorylation) (Kanehisa and Goto 2000, Kanehisa 2019, Kanehisa, Furumichi et al. 2021) showed three types of transient expression dynamics; early upregulation (FB4-6), intermediate upregulation (FB6-8) and late upregulation (FB9-12) (Figure 4j-l). In FB4-6, both CMS36 and CMS30 did not show *CiINV* upregulation, whereas *CiVI* was upregulated together with genes involved in glycolysis in CMS36 and CMS30 (Figure 4j). The lack of *CiINV* upregulation in FB4-6 may suggest that less sucrose is available and perhaps there is insufficient sucrose to support proper functioning of anther tissues and early pollen development in CMS clones. In mid stages (FB6-8), genes involved in glycolysis showed weak upregulation in CMS36, and no upregulation in CMS30 (Figure 4j). In late stages (FB9-12), CMS36 displayed weak upregulation of *CiINV* and *CiVI*, while CMS30 displayed no upregulation (Figure 4i). Accordingly, microscopic analysis revealed that CMS36 pollen are degrading and CMS30 contained no pollen at FB9-12, suggesting that *CiINV* and *CiVI* expression was needed for carbohydrate accumulation during late stages of pollen development (Figure 3n-p,v-x).

### Anther dehiscence

In the final stage of anther development, anther dehiscence releases mature pollen from the anther. This is a complex process that involves coordination of pollen maturation and degradation of specialized cell types in the anther, and defects in timing or failure of dehiscence results in male sterility (as reviewed by Wilson, Song et al. (2011)). Anther dehiscence requires three events: expansion and lignification of the endothecium; septum degradation to form a bi-locular anther; and finally stomium degradation for anther opening (Goldberg, Beals et al. 1993). Microscopic analysis of fertile flower buds revealed that septum degradation started at FB7 and finished at FB12, endothecium lignification took place at FB10, and stomium degradation resulting in anther opening occurs at FB12 (Figure 3d-h). We identified a specific cluster with about 20 co-expressed genes that showed a peak at FB8, followed by temporal suppression at FB9, and subsequent upregulation in FB10 and FB12 (Figure 4m). Two lines of evidence suggest that this cluster may reflect a functional module that regulates anther dehiscence. First, the presence of jasmonate response genes in the cluster suggests that expression of these genes might be regulated by jasmonate, a known regulator of anther dehiscence. Second, the cluster also contained genes involved in cell wall degradation and modification such as polygalacturonases, expansin, cellulase, pectinase and invertase. Such cell wall degrading enzymes are involved in septum and stomium degradation (Wilson, Song et al. 2011). Likewise, a class III peroxidase was previously associated to lignification of the endothecium cell wall (Shigeto, Itoh et al. 2015).

Microscopic analysis of CMS flower buds revealed that anther opening occurs around FB8 (CMS36) and FB9 (CMS30), thus earlier compared to the fertile line (FB12, Figure 3h,m,v). This was accompanied by a temporal shift in endothecium lignification, which occurred at FB6 in both CMS clones, earlier compared to the fertile line, where endothecium lignification occurs at FB9 (Figure 3f,k,s). Start of septum degradation was not observed in CMS clones, but the septum was fully degraded by the time of anther opening (Figure 3m,v). We also found two marker genes for dPCD, which were upregulated at FB9-10 in the fertile line; *BIFUNCTIONAL NUCLEASE1* (*CiBFN1*) and *RIBONUCLEASE3* (*CiRNS3*) (Olvera-Carrillo, Van Bel et al. 2015). In CMS clones, upregulation of these genes started at FB7, indicating that dPCD processes also drive the earlier timing of anther dehiscence in CMS clones (Figure 4n).

Notably, the jasmonate-regulated gene cluster showed different expression dynamics in both CMS clones compared to the fertile line, as most genes were progressively upregulated starting at FB9 until FB12, lacking the transient FB8 peak in the fertile line (Figure 4m). In CMS30, anther dehiscence was also characterized by an inwards collapse of the anther (Figure 3v). Endothecium lignification, septum degradation, anther dehydration, and pollen swelling are all necessary for correct anther opening, and especially dehydration of endothecium and epidermal cells are required for outward bending of the anther (Wilson, Song et al. 2011). In both barley and rice, pollen swelling is partially responsible for the force for rupture of the partially degraded septum (Matsui, Omasa et al. 2000, Matsui, Omasa et al. 2000). Additionally, pollen swelling might result from water relocation because of an increase in potassium ions in the pollen, as demonstrated in multiple species (Bashe and Mascarenhas 1984, Heslop-Harrison, Heslop-Harrison et al. 1987, Matsui, Omasa et al. 2000). It is possible that the increase in potassium ions in the pollen attracts water from the surrounding tissues and might contribute to the dehydration of the endothecium and epidermis (Rehman and Yun 2006, Wilson, Song et al. 2011). Conversely, the absence of intact pollen in CMS30 could thus explain the inward collapse of the anther (Figure 3t-u).

### Plant hormones

To investigate the regulatory role of plant hormones during flower bud development, we analyzed the expression profiles of genes involved in biosynthesis and response pathways of abscisic acid, auxin, brassinosteroids, cytokinins, ethylene, gibberellin and jasmonate (Kanehisa and Goto 2000, Nemhauser, Hong et al. 2006, Kanehisa 2019, Kanehisa, Furumichi et al. 2021) (see Materials and Methods, Figure 5 and Supplemental Data Set 2). We found putative response genes for most hormones upregulated in early, mid and/or late stages (Figure 5). Here, we discuss the most noteworthy expression profiles based on co-expression of rate-limiting phytohormone biosynthesis enzymes and response genes, contrast a fertile line with CMS clones, and place them in the frame of the known role of hormones in anther development as previously described in literature.

**Figure 5.**
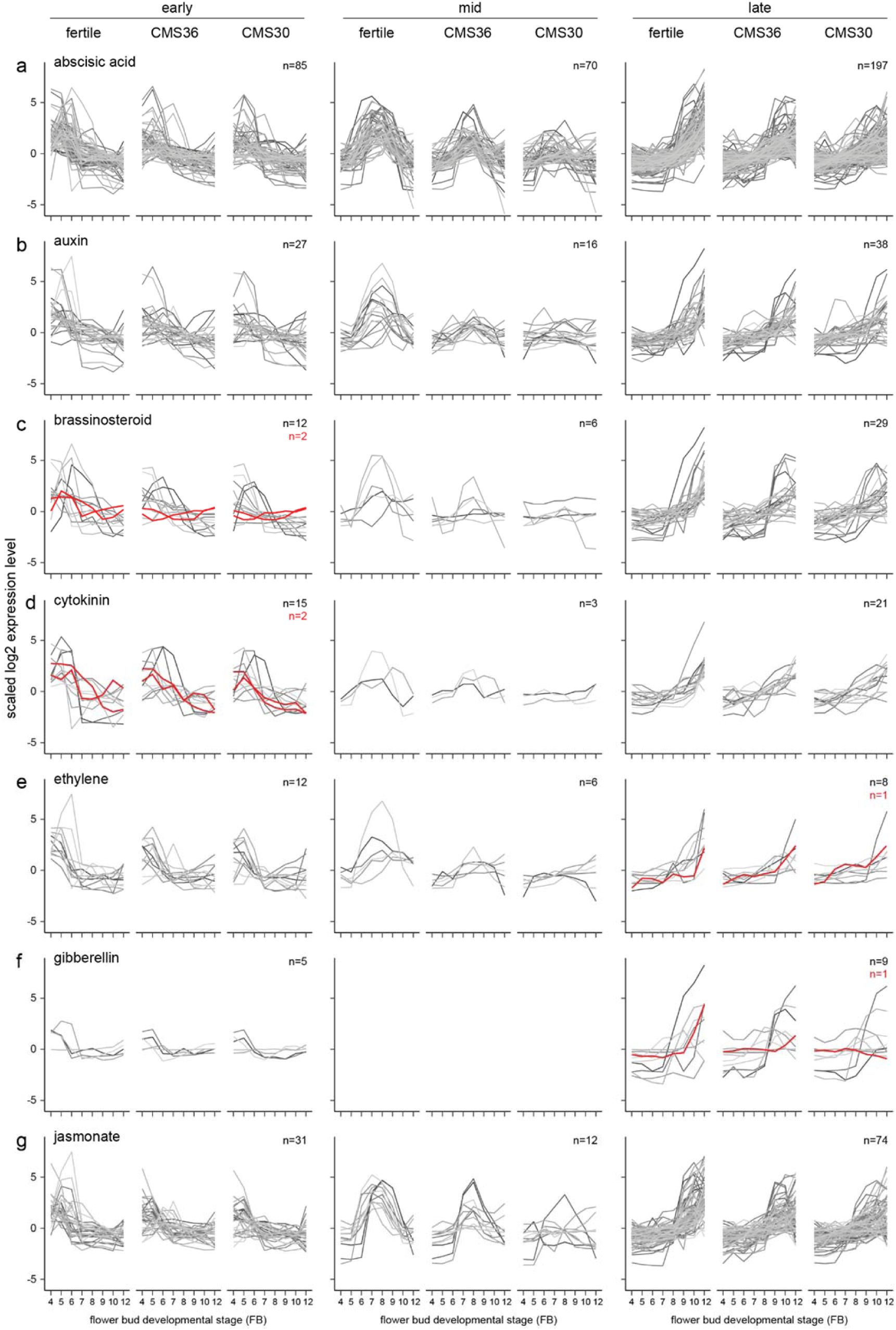
Expression profiles of phytohormone biosynthesis pathway genes (red) and response genes (greyscale) of fertile line K1337 and male sterile CMS clones CMS36 and CMS30 of industrial chicory (*Cichorium intybus* var. *sativum*). x-axis shows flower bud developmental stages, ranging from 4mm (FB4) to 12mm (FB12). y-axis shows scaled log2 Reads Per Kilobase transcript per Million reads mapped (RPKM) values. Gene identifiers per subgroup are listed in Supplemental Data Set 2. For information on ortholog assignment, see Materials and Methods.

### Early FB stages

In early stages of flower bud development (FB4-6) two rate-limiting brassinosteroid genes (*BRASSINOSTEROID-6-OXIDASE)* are upregulated. In CMS clones, these genes are not upregulated, similar to observations of *CiINV* in early stages (Figure 5c, 4i). In rice, brassinosteroid signaling is important for establishing sink strength of the developing anther and pollen grains, and defects in brassinosteroid synthesis/signaling led to reduced starch accumulation in pollen grains (Zhu, Liang et al. 2015), indicating that brassinosteroids could be linked to the energy metabolism in chicory.

### Intermediate FB stages

In the intermediate stages of flower bud development (FB6-9) in the fertile line, auxin response genes are highly upregulated, suggesting transient activity of the auxin-dependent pathway (Figure 5b, mid). In contrast to the fertile line, transient auxin response gene expression is partially suppressed in CMS36, and almost completely suppressed in CMS30 (Figure 5b, mid). Furthermore, microscopic analysis showed that CMS36 anthers contain intact pollen, while CMS30 anthers contain degrading pollen at stages FB6-8 (Figure 3k-m,s-u). As developing pollen accumulate auxin (Cecchetti, Altamura et al. 2008, Salinas-Grenet, Herrera-Vasquez et al. 2018), our observations suggest that absence of normal pollen development explains, at least in part, the lack of upregulation of genes involved in pollen development in CMS clones, as well as auxin-dependent gene expression. In addition, a limited number of ethylene response genes were upregulated during FB6-9, and showed a similar expression profile to auxin response genes in all three genotypes (Figure 5e, mid). Co-expression of genes involved in auxin, ethylene, and brassinosteroid metabolism, response, and/or signaling thus potentially facilitates hormonal crosstalk in the developmental stage after tapetum degradation and before endothecium lignification (FB6-9).

### Late FB stages

In late stages (FB9-12) of flower bud development in the fertile line and CMS clones, jasmonate response genes are upregulated together with ethylene and gibberellin biosynthesis and response genes, suggesting transient activity (Figure 5e-g, late). Late jasmonate response genes include genes involved in programmed cell death, cell wall modification, anther dehiscence, carbohydrate transport, and auxin and gibberellin response. Both ethylene and gibberellin response genes include cell wall modification, and gibberellin response genes also include jasmonate biosynthesis and pollen development genes. These results suggest that these hormones regulate anther dehiscence, as cell wall modification and programmed cell death are key processes for anther dehiscence (Wilson, Song et al. 2011). Genes linked to auxin response in late stages were also involved in cell wall modification, gibberellin and ethylene pathways. Taken together, these results suggest that concerted action of jasmonate, gibberellin, ethylene, and auxin regulate anther dehiscence in chicory.

## DISCUSSION

### Genomic resource to build a comprehensive overview of gene regulatory networks controlling flower bud development

In this study, we report a new genome assembly for industrial chicory (*C. intybus* var. *sativum*), complete with structural annotations of genes. Furthermore, we identified genes involved in flower bud development and CMS, which were assigned to biochemical pathways and were manually curated. The genome, as well as all predicted and curated genes are available on ORCAE as a public resource and can be used for further genetic studies, comparative genomics and analysis of chromosome structural elements.

### Fertile anther development is strictly timed and regulated

Based on the timing of morphological processes observed via light microscopy, combined with the temporal expression profiles of genes and their functional annotation, we were able to reconstruct the temporal sequence of morphological, cellular, and metabolic events, and link those to hormonal action and regulatory genetic pathways. Using the contrast in gene expression profiles between the fertile line and male sterile CMS clones, we could assign some of these pathways to anther and pollen development, and reconstruct a timetable of molecular events (Figure 6). In this discussion, the processes described in the results will be integrated and placed in their respective tissue context during fertile flower bud development: the tapetum as feeding tissue supporting pollen development; pollen development; and the role of epidermis, endothecium and phytohormones in anther opening (Figure 6).

**Figure 6.**
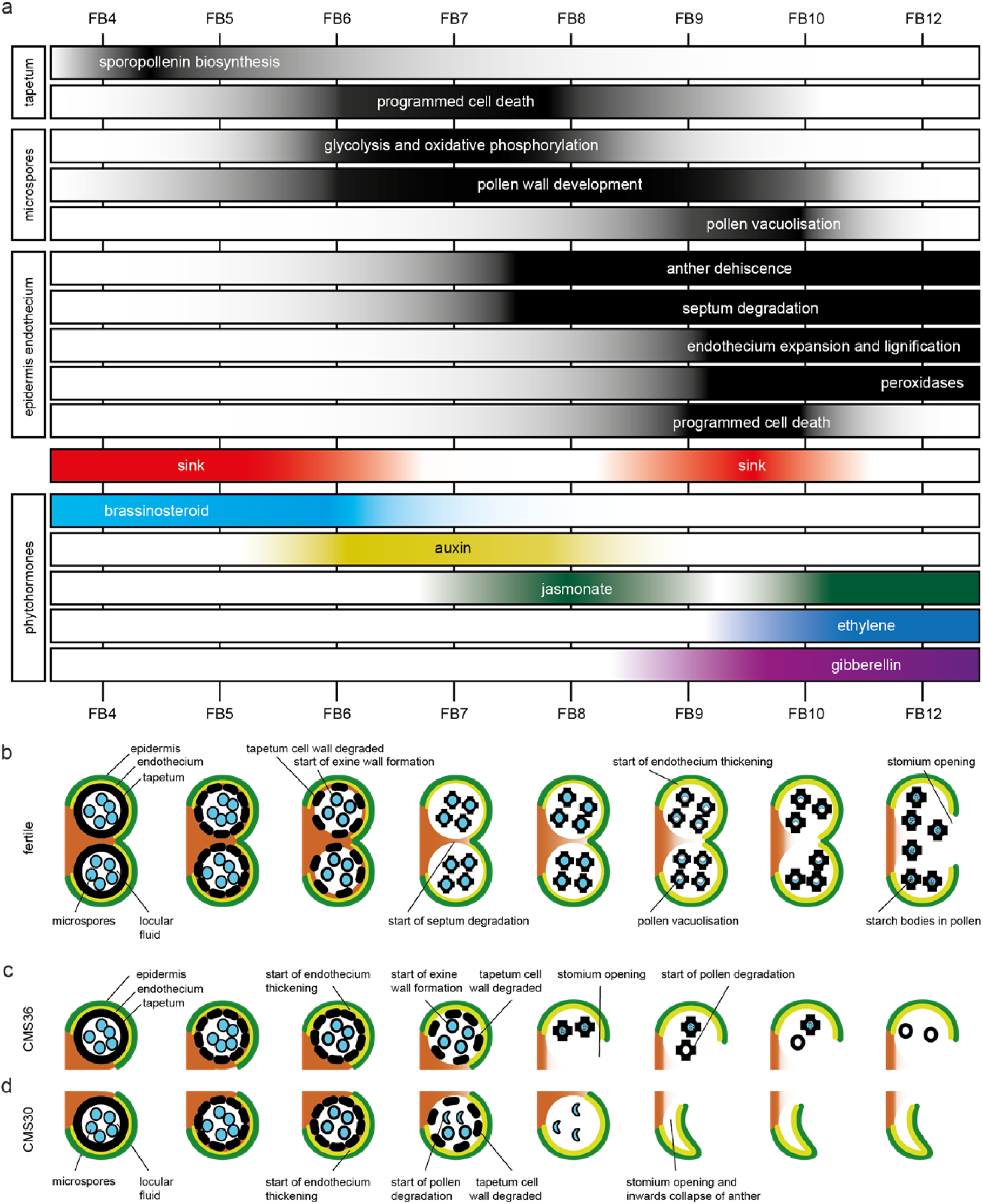
Timetable of molecular, cellular, metabolic and morphological events during anther development in fertile and CMS clones of industrial chicory (*Cichorium intybus* var. *sativum*). a) Overview of transient activity of processes in the tapetum, developing pollen, epidermis, endothecium, and activation of sink strength and phytohormones. b) schematic overview of fertile anther development in chicory. c) schematic overview of anther development in male sterile chicory clone CMS36. d) schematic overview of anther development in male sterile chicory clone CMS30.

### Tapetum as feeding tissue

In the fertile line, the tapetum is present during FB4-5. During these stages, sporopollenin biosynthesis genes, as well as important transcriptional regulators (including *CiTDF1*, *CiMS188*, *CiTEK*, and *CiMS1*) are highly expressed (Figure 6a). In turn, expression of *CiVI* and *CiINV* facilitate that the anther acts as a strong sink tissue early during flower bud development (FB4-6) and help fulfill the high metabolic needs of the tapetum (Figure 6a). During these stages, brassinosteroid biosynthesis genes were upregulated early during normal pollen development but not in CMS clones, and are potentially linked to the sink strength of the anther (Figure 6a) (Zhu, Liang et al. 2015). From FB6 until FB9, dPCD marker genes (aspartic proteases) are upregulated and indicate breakdown of the tapetum to release pollen wall compounds and sugars for the developing pollen in the locule (Figure 6a).

### Pollen development

At early stages (FB4-5), tapetum cell walls are intact and developing pollen do not have a pollen wall yet, which starts to develop around the time of tapetum cell wall degradation (FB6) (Figure 6b). The pollen wall develops further until after the onset of anther dehiscence (Figure 6b). Exine wall formation during FB6-8 also coincides with HCAA biosynthesis, which are important components of the pollen wall. During FB6-9, genes involved in glycolysis are upregulated, which could be a signature of pollen metabolism (consistent with absence of upregulation of these genes in CMS clones that lack active pollen) (Figure 6a). In addition, both the MVA and MEP pathway for isoprenoid synthesis were active in early FB stages. The isoprenoid biosynthesis pathway produces the precursors of many important compounds, such as certain plant hormones, but also sterols and terpenes (Vranova, Coman et al. 2013, Tholl 2015). Two hormones may be associated with the development and metabolic activity of pollen. Auxin response genes showed co-expression with energy metabolism genes in both the fertile line and CMS clones, coinciding with early stages of pollen development, and consistent with previous reports that developing pollen accumulate auxin (Cecchetti, Altamura et al. 2008, Salinas-Grenet, Herrera-Vasquez et al. 2018) (Figure 6a). In FB9-10 fertile pollen start to vacuolize, which coincides with a second period of upregulated *CiVI* and *CiINV* expression, suggesting that the anther again acts as a sink, and pollen accumulate sucrose to establish a carbohydrate pool (Figure 6a-b). In addition, the upregulation of *CiVI* also suggests that pollen further convert sucrose to glucose and fructose, which creates an osmotic effect needed for pollen vacuolization.

### Role of epidermis, endothecium and phytohormones in anther opening

During the last stages of anther development, morphological changes in endothecium and epidermis drive anther dehiscence. In the fertile line, this starts at FB7 with the onset of septum degradation (Figure 6b). At FB9, the endothecium starts expanding and endothecium cell walls become lignified, consistent with subsequent upregulation of genes involved in monolignol biosynthesis (FB6-9) and monolignol polymerization by peroxidases (FB10-12) (Figure 6a-b). Finally, stomium degradation results in anther opening and release of mature pollen.

Several lines of evidence suggest that auxin, jasmonate, ethylene, and gibberellin interact to regulate the timing of anther dehiscence events. As described above, in FB6 auxin response genes are transiently expressed in developing pollen. In *Arabidopsis,* auxin accumulation has been linked to synchronization of pollen development and anther dehiscence, such as the timing of the lignification process that marks the start of anther dehiscence (Cecchetti, Altamura et al. 2008, Salinas-Grenet, Herrera-Vasquez et al. 2018).

The transient increase in auxin response genes is followed by a sharp and transient peak in jasmonate response genes at FB8 (Figure 6a) during fertile flower bud development. After a marked transient downregulation of jasmonate-responsive gene expression, the jasmonate-dependent genes are again upregulated in FB10-12 (Figure 6a). As previously described, jasmonate is known to be involved in anther dehiscence in many species, as evidenced by indehiscent anthers of jasmonate biosynthesis mutants (Wilson, Song et al. 2011), and exogenously applied high jasmonate concentrations induce precocious anther dehiscence (Cecchetti, Altamura et al. 2013). In addition, auxin can negatively regulate jasmonate biosynthesis and such hormonal crosstalk may regulate the timing of anther dehiscence (Cecchetti, Altamura et al. 2013). Therefore, we postulate that the transient peak of jasmonate response genes at FB8, potentially in combination with negative regulation by auxin, has an important regulatory role in the timing of anther dehiscence.

Together with upregulation of jasmonate responsive genes in FB10-12, ethylene-dependent gene expression in FB10-12 was observed, while gibberellin response genes are upregulated during F9-12, suggesting hormonal crosstalk between these three phytohormones (Figure 6a). Gibberellin is important for stamen development, and is involved in filament elongation, tapetum development, pollen formation, and anther dehiscence in *Arabidopsis* (Wilson, Song et al. 2011). Both gibberellin and jasmonate are needed for anther dehiscence in *Arabidopsis* and crosstalk between gibberellin and jasmonate-dependent pathways is required for stamen development (Peng 2009, Marciniak and Przedniczek 2019). A role for ethylene during anther dehiscence has been shown in petunia and tobacco (Rieu, Wolters-Arts et al. 2003, Wang and Kumar 2007). During FB10-12, also extracellular peroxidases are upregulated, known to be important for monolignol polymerization (Figure 6a).

### Energy deficiency and retrograde signaling models may explain male sterility in CMS36 and CMS30

Global analysis of the FC4 gene set revealed that gradually more processes become uncoupled in CMS compared to the fertile line, as evidenced by PCA analysis, and the fraction of divergent gene expression profiles in CMS clones compared to the fertile line (Figure 2). Here, we provide an overview of processes that are affected in CMS clones, and evaluate the four possible mechanisms underlying CMS.

In the tapetum of CMS36 and CMS30, present during stages FB4-6, both the genetic regulators and sporopollenin genes were upregulated in early stages, but display slightly delayed developmental timing, which was consistent with microscopic observations of delayed tapetum degradation (Figure 4a, Figure 6b-d).

The pollen in CMS36 develop normally during FB4-8 but start to degrade after anther opening, while the pollen in CMS30 initially develop normally, but start to degrade at FB7 (Figure 6b-d). Glycolysis, oxidative phosphorylation, and auxin-response genes are less (CMS36) or not (CMS30) upregulated during stages FB6-9, indicating that auxin accumulation is not taking place in CMS clones. Likewise, brassinosteroid genes are not upregulated in CMS clones. The lack of upregulation of brassinosteroid-associated genes, in turn, could be explained by the suppressed expression of isoprenoid and sterol biosynthesis pathways in CMS clones, which normally produce precursors for brassinosteroid biosynthesis.

Anther opening is premature in CMS36 (FB8) and CMS30 (FB9) compared to the fertile line (FB12), which could be due to disturbed phytohormone regulation (Figure 6b-d). Auxin accumulation is important for the timing of anther dehiscence processes, but is decreased in developing CMS pollen. It is possible that the altered auxin levels at stages FB6-9 further lead to the disturbed jasmonate signaling in later stages, and subsequently earlier timing of dehiscence. These gene expression dynamics suggest that deregulated jasmonate induced processes, together with lack of auxin-jasmonate crosstalk, could explain the earlier timing of anther opening in CMS36 and CMS30.

Based on these results, we can evaluate which CMS model fits our observations best. As tapetum degradation seems to occur similarly in the fertile line and CMS clones, despite a slight delay in timing, evidenced by microscopy and expression data from dPCD markers, the aberrant PCD model is an unlikely mechanism. The cytoxicity model could explain the degradation of developing pollen in CMS30 and for CMS36 in later stages. In line with the energy deficiency model, a lower sink strength in early and late stages (as evidenced by gene expression of *CiINV*), may suggest that tapetum energy requirements to support pollen development may not be met. As both CMS36 and CMS30 were established by crossing a wild chicory genotype (as mother) with an industrial chicory inbred line (as father), followed by subsequent backcrosses, it is likely that the male sterility phenotype is the result of incompatibility between mitochondria of the wild chicory and nuclear genome of the industrial chicory, compliant with the retrograde regulation model. In summary, our results are mostly in line with the energy deficiency model, and/or disturbance of retrograde signaling from mitochondrion to nucleus, or a combination of multiple mechanisms (Chen and Liu 2014).

## Conclusion

In conclusion, we provided a new industrial chicory reference genome assembly with an annotated and curated gene set related to flower bud development and male sterility that can be used for further genetic studies. The genome can also be used as a resource for genetic studies related to processes not discussed in this paper. Additionally, we provided a comprehensive molecular and morphological overview of processes related to chicory anther development and processes which are disturbed in CMS. This overview can be the basis of further studies on anther development and male sterility in chicory.

## METHODS

### Plant material

For genome sequence assembly, an eighth generation inbred line of industrial chicory (*C. intybus* var. *sativum*, L8001; derived from family H79 from the ILVO chicory breeding program), was used for genomic DNA extraction (see below). For transcriptome and morphological analysis of flower buds, an industrial chicory clone (K1337) and an inbred line (L8018) were used as fertile control lines with normal pollen production (referred to as ‘fertile’), in addition to CMS clones 12-712-30 (CMS30) and 12-712-36 (CMS36). Both CMS clones can become fertile under favorable environmental circumstances, such as elevated temperatures during anther development, but CMS30 shows more stable male sterility, while CMS36 shows full fertility restoration (Van der Veken, Vandermoere et al. 2018). All plants were kept on the field protected by a plastic cover during winter for vernalization and grown on a container field until bolting. Upon bolting, plants were transferred to a controlled growth chamber at 16h/8h day/night with temperature of 15 ± 2°C and SONT lights (MASTER GreenPower 600W EL Plus, Philips, The Netherlands) with a photosynthetic photon flux density (PPFD) of 210 μmol.m^-2^.s^-1^ measured at 1 m above the floor using a LI-250A light meter (LI-COR, United States). Various tissues were sampled from line L10003 (two further generations of selfing after L8001), grown under similar conditions, to create the small tissue panel.

### DNA extraction for genome assembly

For PacBio and Illumina whole genome shotgun (WGS) sequencing, high molecular weight (HMW) genomic DNA was extracted with a modified CTAB protocol (Murray and Thompson 1980). In short, 0.5 g of chicory root powder was incubated in 2% CTAB buffer at pH 5.0, supplemented with 0.2% β-mercaptoethanol and 1% PVP-40, for 1 hour at 60°C with regular inversion. The mixture was extracted twice with chloroform:isoamylalcohol (24:1), and precipitated by adding 0.5 volume of 5 M NaCl, 0.05 volume 7.5 M ammonium acetate and 0.6 final volume ice-cold isopropanol. After centrifugation (15 min, 4°C, 4000 rpm), the pellet was washed three times with 70% ethanol. The pellet was resuspended in 10 mM Tris pH 8.0 and treated with RNAse A, followed by phenol:chloroform:isoamylalcohol (25:24:1) extraction, chloroform:isoalylalcohol extraction and ammonium acetate/ethanol precipitation. The pellet was finally resuspended in 10 mM Tris pH 8.0.

Several approaches for extracting HMW nuclear genomic DNA were tried, but the best performing protocol for ONT sequencing was similar to the BioNano Prep High Polysaccharides Plant Tissue DNA Isolation protocol (document number 30218, revision C, www.bionanogenomics.com). In short, etiolated fresh young leaves of industrial chicory line L8001 were chopped into a fine mesh with a razor blade. This material was filtered to recover nuclei, which were embedded into CleanCut Low Melting agarose plugs (Bio-Rad). Five plugs per falcon tube were incubated in BioNano Prep Lysis Buffer (BioNano Prep Plant DNA isolation kit; part # 80003), with proteinase-K (Qiagen) at 50°C overnight with intermittent mild shaking. The next day, the proteinase K solution was refreshed and incubated for another 2 h at 50°C, followed by RNase A (Qiagen) treatment at 37°C for 1 h. Subsequently, the agarose plugs were washed (5 times 15 min) and then melted at 70°C for 2 min in an 1.5 ml Eppendorf tube (1 plug per tube) placed at 43°C, and agarase (ThermoFisher Scientific) and RNase A were added and incubated at 43°C for 1 h. The material from the melted and digested plug was then dialyzed over a 0.1 µm dialysis membrane (Millipore) floating on a TE wash buffer for 2 h. The remaining drop on the membrane was used as the starting material for library construction for ONT sequencing (HMW genomic DNA).

For HiC sequencing, nuclei were first extracted from chicory leaf protoplasts. Protoplasts were obtained after 3 hours incubation in digestion buffer (1.5% cellulose R10, 0.4% macerozyme, 0.5M mannitol, 20mM KCl, 20mM MES pH5.5, 10mM CaCl_2_). Debris was removed by filtering the digestion mixture using a 100µM filter, followed by centrifugation (100g, 10 min, 20°C). Protoplast were further purified using a sucrose gradient. Protoplast were resuspended in nuclei isolation solution pH 7 (0.1mM spermidine, 10mM phosphate buffer pH 7, 2.5mM EDTA, 10mM NaCl, 10mM KCl, 0.2M sucrose, 0.15% Triton X-100, 2.5mM DTT), and formaldehyde was added (1% final concentration), followed by 15 minutes incubation at RT. Glycine was added (0.2M final concentration), followed by 20 minutes incubation at RT and subsequently 7 minutes on ice. Protoplasts were transferred to large syringes, and passed 4 times through a 25G 5/8 gauge needle. The lysate was subsequently filtered through a 10µM filter, followed by centrifugation (400g, 10 min). The pellet was resuspended in another nuclei isolation buffer (0.1mM spermidine, 10mM MES-KOH pH5.5, 2.5mM EDTA, 10mM NaCl, 10mM KCl, 0.2M sucrose, 0.15% Triton X-100, 2.5mM DTT). The lysate was again filtered (10µM filter), centrifuged (400g, 10 min) and stored in 40% glycerol. Hi-C libraries using DpnII were generated according to Liu, Cheng et al. (2017).

### Sequencing and assembly

The PacBio data was generated on an RS II system, using P6 chemistry (Wageningen University, NL). The Illumina data was obtained with Illumina HiSeq2500 (WGS and Hi-C), MiSeq, and NextSeq500 (KULeuven Genomics Core facility, Belgium). ONT sequencing was done using the MinION Mk1B, where the Ligation Sequencing Kit 1D (SQK-LSK108, ONT) and Rapid Sequencing Kit (SQK-RAD0004, Life Technologies) were used for library preparation and tested on a FLO-MIN106 flow cell. Finally, a library was built from the obtained genomic DNA using the ligation protocol (SQK-LSK108) and sequenced (72h run) on a fresh FLO-MIN106 flow cell. All the results from the sequencing tests and the final full run were combined for assembly. Base calling of the ONT raw data was done with guppy v3.4.1. The data were primarily assembled using the longer ONT reads combined with PacBio reads, while the Illumina reads contributed to the polishing of the genome. The ONT reads were assembled using the Flye software (v2.5), with --min-overlap 3000 to increase stringency at the initial overlay step, -ont-raw to accommodate the ONT error rate, and default parameters including nine rounds of polishing through consensus (Kolmogorov, Yuan et al. 2019). Long-Read scaffolding was attempted with the PacBio reads. This assembly was further polished using Pilon (Walker, Abeel et al. 2014), with Illumina short reads. The assembly was purged from haplotigs and repeat-containing contigs using purge_haplotigs. The Hi-C reads were used to scaffold the 13490 contigs into 10515 Hi-C scaffolds using Juicer (Durand, Shamim et al. 2016) and the 3D-DNA (v180114) pipeline in diploid mode, with two rounds of mis-join correction. Contigs <15 kb were not considered at this stage to reduce ambiguous mapping of the Hi-C read pairs. At this stage, the total assembly of 948 Mb contained 17 large scaffolds (N90 = 40 Mb) roughly representing chromosome arms, together with a large set of shorter contigs. These Hi-C scaffolds were further ordered and linked into the nine chromosomes using a genetic map. The resulting assembly yielded an anchored genome of 877 Mb. A final polish was applied using Medaka (https://github.com/nanoporetech/medaka) (v0.11.2, ONT data) and Racon (Vaser, Sovic et al. 2017) (PacBio data) aiming at filling gaps resulting from the Hi-C and genetic map scaffolding. BUSCO genome completeness analysis was performed using the eudicots_odb10 reference set of 2326 genes (Manni, Berkeley et al. 2021). Whole genome triplication events were analyzed with i-ADHoRe software (Proost, Fostier et al. 2012).

### Gene prediction

Protein coding genes were predicted using Augustus (Stanke, Diekhans et al. 2008) on the polished and frozen reference genome sequence, including hints from RNA-Seq (flower development and tissue panel) and alignments with proteins from *Arabidopsis,* tomato and lettuce retrieved from diverse publicly available sources. To reduce the degree of gene model over-prediction due to repeated elements (transposable elements and simple sequence repeats), high abundant repeats were *de novo* predicted with RepeatModeler (https://github.com/Dfam-consortium/RepeatModeler). The obtained repeat library for chicory was used to soft-mask the genome sequence with RepeatMasker. Further curation of the predicted gene models was done via the ORCAE interface (Sterck, Billiau et al. 2012). BUSCO gene set completeness analysis was performed using the eudicots_odb10 reference set of 2326 genes.

### Sampling of flower bud developmental stages

For transcriptome analysis, floral buds of the fertile line K1337 and CMS clones CMS30 and CMS36 were sampled in the growth chambers between 9:30 and 11:30 AM. Buds were divided into eight size categories using a digital caliper to measure bud size: 3-4 mm (FB4), 4-5 mm (FB5), 5-6 mm (FB6), 6-7 mm (FB7), 7-8 mm (FB8), 8-9 mm (FB9), 9-10 mm (FB10) and 10-12 mm (FB12) (see Figure 6). At least five floral buds of three different plants of the same clone were pooled. For the samples containing the smallest bud sizes, up to 20 floral buds were harvested to obtain sufficient material for RNA extraction. Samples were immediately frozen in liquid nitrogen and stored at -80°C until RNA extraction. Two biological replicate samples were taken per clone per size category. For morphological analysis, samples of fertile line L8018, CMS30 and CMS36 were harvested in identical conditions and size categories representing floral bud developmental stages (FB4-FB12).

### RNA extraction and sequencing

Pooled samples were ground and 100 mg tissue was used for RNA extraction. Total RNA was extracted using a modified CTAB protocol (Luypaert, Witters et al. 2017), including DNase treatment (DNA-free, Invitrogen). RNA quantity and integrity were measured using Nanodrop ND-1000 (Thermo Fisher) and Experion (Biorad) instruments, respectively. For all samples, 3 μg (50 ng μL^-1^) of RNA was used for TruSeq stranded RNA-Seq library preparation, quality control with Agilent Technologies 2200 TapeStation and paired-end (2x100 bp) sequencing on an Illumina NovaSeq6000 instrument (Macrogen, Korea).

### RNA-Seq data analysis

RNA-Seq reads were trimmed, de-replicated and mapped against the CDS sequence of 54020 predicted genes of the novel *C. intybus* var. *sativum* reference genome sequence (https://bioinformatics.psb.ugent.be/orcae/overview/Cicin), using BWA-mem (Li and Durbin 2009). Data analysis was done using R v.4.0.5 with the package edgeR (McCarthy, Chen et al. 2012). Genes with expression in less than two samples and with less than four Reads Per Kilobase of transcript per Million mapped reads (RPKM) were discarded. Normalization to scale the raw library sizes was done using the trimmed mean of M-values (to the reference, selected using default settings) (Robinson and Oshlack 2010). A principal component analysis (PCA) was performed with log2 transformed counts of all samples using the DESeq2 package in R v4.0.5 (Love, Huber et al. 2014). To identify differentially expressed genes (DEGs) of the fertile line (K1337) between the eight developmental stages FB4-FB12, we fitted a negative binomial generalized log-linear model to the RPKM values for each gene. We identified lowest and highest expression level within the fertile line developmental series and subsequently performed likelihood ratio tests (LRT). Thresholds for further filtering of the DEGs were set at minimal 4-fold-change in expression (log2-fold-change of 2, here called the FC4 gene set) and an adjusted p-value (Benjamini-Hochberg multiple testing correction) cutoff of 0.05 (Benjamini and Hochberg 1995).

Differentially expressed genes were assigned to co-expression clusters based on the developmental stage of their maximum expression level. Clustering was performed using the Mfuzz package in R v.4.0.5 (Kumar and E Futschik 2007), followed by manual curation. Genes with different expression profile in one or both CMS clones compared to the fertile line were manually selected by examining the expression profiles.

### Gene orthology analysis of FC4 gene set

To assign genes in the FC4 gene set to certain biochemical pathways, *Arabidopsis* gene identifiers were extracted from KEGG (Kanehisa and Goto 2000, Kanehisa 2019, Kanehisa, Furumichi et al. 2021), while *Arabidopsis* genes with a transcriptional response to a given phytohormone were extracted from Nemhauser, Hong et al. (2006). These coding DNA sequences (CDS) were used as queries to identify the *C. intybus* homologs using a tBLASTn search (e-value <1e-50), against the protein sequences of all predicted *C. intybus* gene models. Orthologs that belonged to the FC4 gene set were extracted and their (co-)expression pattern was further analyzed in the fertile line and CMS clones. For the hormone response genes, multiple response genes were not unique for one hormone, but an overview of genes involved in these processes is given in Supplemental Data Set 2. ORCAE was used for manual curation of the structural gene annotation of these genes (Sterck, Billiau et al. 2012). Each predicted gene model was manually curated, using available supporting data in ORCAE (such as RNA-Seq read alignments, splicing sites, gapped alignments of *de novo* assembled transcripts, and blast hits of orthologues from closely related species).

### Phylogenetic analysis of genes in biochemical pathways

To identify the correct ortholog for genes involved in biochemical and genetic pathways (genes described in section “Early tapetum development”, “Phenylpropanoid metabolism”, “Energy metabolism”, “Isoprenoid metabolism”, “Anther dehiscence” and rate-limiting hormone biosynthesis genes in “Plant hormones”) phylogenetic analysis was performed. First, per gene the corresponding homology groups (HOMgroups) with all known orthologues of multiple species (*Arabidopsis thaliana, Lactuca sativa, Daucus carota* and *Helianthus annuus*) were extracted from the PLAZA dicots 5.0 database (https://bioinformatics.psb.ugent.be/plaza) (Van Bel, Silvestri et al. 2021). Next, we performed multiple sequence alignment at the protein sequence level per HOMgroup, followed by construction of a phylogenetic tree (Neighbor-Joining method, Jukes-Cantor model, bootstrap replicates set to 100) in CLC genomics Main Workbench. Gene functions were assigned according to the closest *Arabidopsis* ortholog. For the HOMgroup including extracellular peroxidases, elucidating the true gene function was challenging due to the size of the gene family and high level of protein sequence similarity between different gene family members (Lüthje and Martinez-Cortes 2018). Therefore, it should be noted that we cannot be sure all described extracellular peroxidases are truly involved in lignin polymerization.

### Morphological analysis

Morphological analysis was performed based on the protocol reported by (Habarugira, Hendriks et al. 2015), with modifications. Harvested floral buds were vacuum fixated for 20 min in formalin acetic acid-alcohol (FAA) solution (10:7:2:1 ethanol 99%:dH_2_O:formalin 37%:glacial acetic acid) and then further incubated in FAA for 24 h at room temperature. Fixed samples were stored in 50% ethanol at -20°C until further use. Samples were dehydrated by changing the solvent in two steps (70% and 85% ethanol) to 100% ethanol. Dehydrated samples were prepared for embedding by infiltration with glycol methacrylate (Technovit 7100, Heraeus Kulzer, Germany) in 1.5 mL Eppendorf tubes. Large floral bud samples (>4 mm) were dissected before transfer to the infiltration solution. On initial infiltration with a 1:1 ethanol:Technovit + Technovit with hardener 1 solution, another 20 min vacuum step was performed to enhance infiltration. Afterwards, samples remained for 2 h at 4°C in the 1:1 infiltration solution. Samples were transferred into 100% Technovit solution and incubated overnight. For embedding, 1 mL Technovit hardener 2 was added to 15 mL Technovit solution and samples were embedded. Samples were kept for 1 h at room temperature and were later incubated at 37°C overnight. Next, blocks were sectioned at 5 μm using a HM360 Microtome (Thermo Scientific). Slides were stained for 10 min with 1% Toluidine Blue O (Acros Organics, Geel) in demineralized water and rinsed twice in demineralized water. Slides were subsequently covered and sealed with DPX new mounting medium (Merck KGaA, Darmstadt, Germany) and incubated for 3 days prior to microscopic analysis. Slides were analyzed using an inverted microscope with a magnification of 400x and 1000x (Leica DMi8, Wetzlar, Germany) equipped with a camera (Leica DFC450, Wetzlar, Germany) and processed using LAS software (Leica Wetzlar, Germany).

## Accession numbers

All sequencing data was submitted to NCBI SRA under the BioProject number: PRJNA899436; Illumina Short read data: SRR22243842-SRR22243848; Hi-C: SRR22542269, SRR22542268 (restriction site: DpnII); PacBio: SRR22404031; ONT: SRR22704481-SRR22704487). Raw sequence reads of the flower bud development series and the small tissue panel were submitted to NCBI SRA under BioProject PRJNA898887, SRR22213225-SRR22213284.

The genome is publicly available on ORCAE (https://bioinformatics.psb.ugent.be/orcae/overview/Cicin)

## Supplemental Data Files

Supplemental Table 1: Genome assembly and gene annotation statistics.

Supplemental Data Set 1: Cluster assignment and normalized expression levels of FC4 gene set.

Supplemental Data Set 2: Overview table of genes discussed in more detail, including their functional description, assignment per pathway as described in Figures 2, 4 and 5, and normalized expression values during flower bud development.

## ACKNOWLEDGMENTS

We thank Ellen De Keyser and Laurence Desmet for the sampling and RNA extractions of chicory flower buds, Magali Losschaert and Sophie Carbonelle for preparing and staining microscopy sections and subsequent light microscopy and the ILVO field teams for maintenance of plant material. We also thank Bertrand Vandoorne for constructive feedback. Initial sequencing was supported in the framework of a DGA project (Grant D31-1221) with the Walloon region (DGARNE-Belgium). EW and JVdV were supported by a scholarship from Chicoline, a division of Cosucra Groupe Warcoing S.A., Belgium.

## AUTHOR CONTRIBUTIONS

EW, SR, OM and TR designed the research. EW, SR, JB, ND, ADK, TE, AH, CL, OM, JVdV, KVL and TR performed the research. AH contributed new analytic/computational tools. EW, SR, TE, AH, CL, KVL and TR analyzed data. EW, SR and TR wrote the paper with input from all other authors. All authors read and approved the final version of the manuscript.

